# Cell morphological profiling enables high-throughput screening for PROteolysis TArgeting Chimera (PROTAC) phenotypic signature

**DOI:** 10.1101/2022.01.17.476610

**Authors:** Maria-Anna Trapotsi, Elizabeth Mouchet, Guy Williams, Tiziana Monteverde, Karolina Juhani, Riku Turkki, Filip Miljković, Anton Martinsson, Lewis Mervin, Erik Müllers, Ian Barrett, Ola Engkvist, Andreas Bender, Kevin Moreau

## Abstract

PROTACs (PROteolysis TArgeting Chimeras) use the ubiquitin-proteasome system to degrade a protein of interest for therapeutic benefit. Advances in targeted protein degradation technology have been remarkable with several molecules moving into clinical studies. However, robust routes to assess and better understand the safety risks of PROTACs need to be identified, which is an essential step towards delivering efficacious and safe compounds to patients. In this work, we used Cell Painting, an unbiased high content imaging method, to identify phenotypic signatures of PROTACs. Chemical clustering and model prediction allowed the identification of a mitotoxicity signature that could not be expected by screening the individual PROTAC components. The data highlighted the benefit of unbiased phenotypic methods for identifying toxic signatures and the potential to impact drug design.

**Highlights:**

- Morphological profiling detects various PROTACs’ phenotypic signatures
- Phenotypic signatures can be attributed to diverse biological responses
- Chemical clustering from phenotypic signatures separates on drug selection
- Trained *in-silico* machine learning models to predict PROTACs’ mitochondrial toxicity

## Introduction

PROTACs belong to a category of compounds also referred to as beyond the Rule-of-5 (bRo5) as they do not comply with the Lipinski’s Rule-of-5 (Ro5). The prediction and/or better understanding of the consequences for drug screening are limited by the lack of descriptors and methodologies for robust safety profiling. Hence, there is a need for descriptors tailored or ‘compatible’ with the bRo5 new data modalities (Ermondi et al., 2021; Lipinski et al., 1997). There have been machine learning approaches for the prediction of drug toxicity by using physiochemical descriptors, structural alerts and high throughput imaging data for small molecules (Hemmerich et al., 2020; Zhang et al., 2009; Zhao et al., 2021). However, computational prediction for new modalities is less investigated. As a new therapeutic modality, PROTACs are raising multiple concerns on various aspects such as safety, ADME properties, toxicity, and others (Moreau et al., 2020). A potential approach to profile PROTACs and improve understanding of their safety aspects could be the use of high throughput imaging (HTI) assays, which have become easier to run over recent years. HTI assays have been useful in the better understanding of compounds mode of action (Gustafsdottir et al., 2013; Hofmarcher et al., 2019; Simm et al., 2018; Trapotsi et al., 2020, 2021; Young et al., 2008) but from a practical angle have also been used to predict a wide range of efficacy and safety endpoints (Cox et al., 2020; Martin et al., 2014; Seal et al., 2021). One of the assays that is currently used by academic groups and pharmaceutical companies is the Cell Painting assay (Seal et al., 2021; Trapotsi et al., 2020). Phenotypes from this assay are not obtained with any particular biological point of interest in mind and can be considered as image-based fingerprints of a compound covering a wide range of information.

Here, we report for the first time that the Cell Painting assay can be used as a high throughput imaging assay to profile morphological changes induced by PROTACs. Cell Painting descriptors proved to be sufficient to train models with good predictive performance. We proved that these profiles can be useful in mitochondrial toxicity prediction of PROTACs, highlighting that image-based data can be used in both supervised and unsupervised machine learning approaches and provide information for the safety assessment of compounds such as mitochondrial toxicity, which has been related to attrition of drugs and late-stage market withdrawals (Dykens and Will, 2007).

## Results

### Morphological profiling detected PROTACs activity

The study workflow can be divided into four main steps (Figure 1a). PROTACs were profiled with the Cell Painting assay in U-2 OS cells. A total of 341 PROTACs and 149 non-PROTACs, directed at more than 15 different targets, were profiled. The non-PROTAC compounds include small-molecule compounds, which are inhibitors of the targets that PROTACs are degrading, E3 ligase ligands and reference compounds that have shown mitochondrial toxicity. Following the compounds’ profiling with the Cell Painting assay, morphological features were calculated with CellProfiler. Morphological features were normalised, and a feature selection process was applied. In the final step, the activity of PROTACs on Cell Painting assay was evaluated and PROTACs-Cell Painting features were used as descriptors for training the *in*-silico mitotoxicity models.

**Figure 1:**
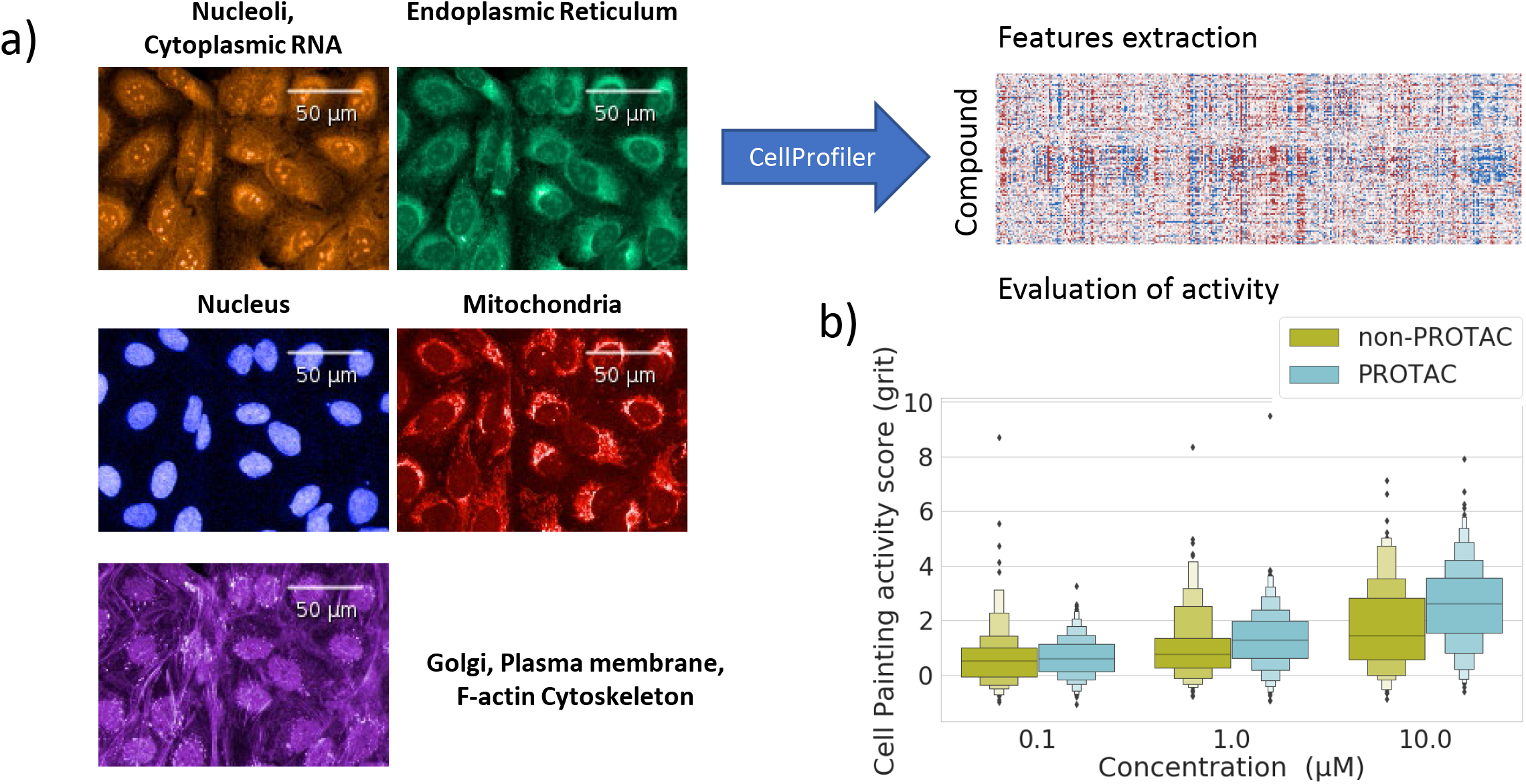
Cell Painting workflow and PROTAC activity. a) Summary of the analysis performed in this work. PROTACs and non-PROTACs compounds were profiled with the Cell Painting assay followed by data normalisation and a downstream analysis. b) Cell Painting activity score in the form of grit score across all concentration (0.1, 1 and 10 μM). Both PROTACs and non-PROTACs compounds’ activity on the cell painting assay (in the form of grit score) increased as the concentration increased.

PROTAC profiles together with non-PROTAC molecules were used to understand whether they show systematically different Cell Profiling readouts compared to neutral controls, based on two metrics: a) Euclidean Distance-based and b) grit score activity metric. The results from the Euclidean distance-based method showed that out of the ~1,000 (3 replicates per PROTAC) profiles obtained from testing PROTACs at concentrations 0.1, 1, and 10 μM, 17%, 61% and 80% of profiles respectively, displayed cellular morphology different from the neutral controls (Figure S1a). In line, higher grit scores were observed with increasing concentration (median ± standard deviation of 0.65±0.72, 1.32±1.07 and 2.56±1.49 for concentrations of 0.1, 1 and 10μM, respectively; Figure 1b). For non-PROTAC compounds, similar trends were observed where 22%, 46% and 60% of a total of ~450 profiles, displayed cellular morphology different from the controls (Figure S1b). Similarly, higher grit scores were observed with increasing concentration (median ± standard deviation of 0.65±1.20, 1.04±1.30 and 1.80±1.60 for concentrations of 0.1, 1 and 10 μM, respectively; Figure 1b). Hence, we observed a clear dose-response relationship in the dataset examined here.

Looking at particular examples, we focused on a commercially available PROTAC dataset, which included PROTACs targeting BRD4 and PROTACs targeting CDK proteins (Table 1 and Figure 2). All previously published PROTACs showed activity in the Cell Painting assay, including PROTACs targeting BRD4 and PROTACs targeting the cell cycle regulators CDK proteins (Figure 2). Among the BRD4 PROTACs, MZ1 and ZXH 3-26 were the most active PROTAC compounds while dBET1 was the least active (Figure 2), matching the degradation potency described for these compounds at BRD4 degradation suggesting that the activity seen is an on-target effect. Among the CDK degraders, the PROTAC targeting CDK9 (THAL-SNS-032) was the most active. This makes it a pharmacologically interesting PROTAC because of its selective degradation of CDK9 with limited effects on the protein level of other CDKs (Olson et al., 2017). In addition, THAL-SNS-032 has shown a prolonged pharmacodynamic effect compared with traditional kinase inhibitors (Olson et al., 2017). Looking at the raw images, it was clear that the CDK9 degrader caused a reduction in nucleoli formation, suggesting a cell cycle arrest effect, in line with the function of CDK9 in cell cycle progression (Figure 2). This phenotype is plausible given that CDK9 inhibitors - such as the Flavopiridol – promote nucleolar disintegration by inhibiting early rRNA processing and transcription (Carotenuto et al., 2019).

**Table 1:**
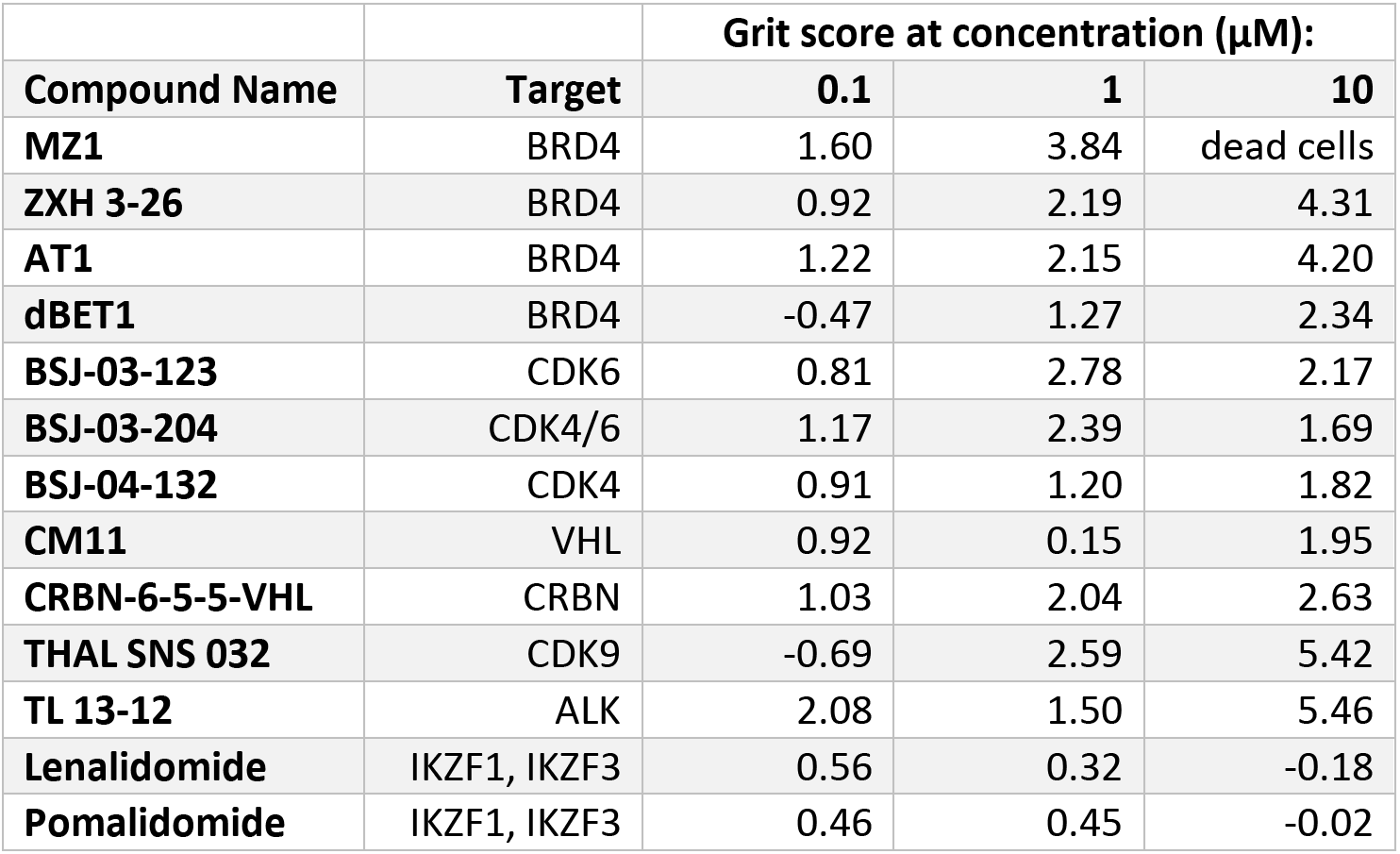
Cell Painting activity score (Grit) for published PROTACs

**Figure 2:**
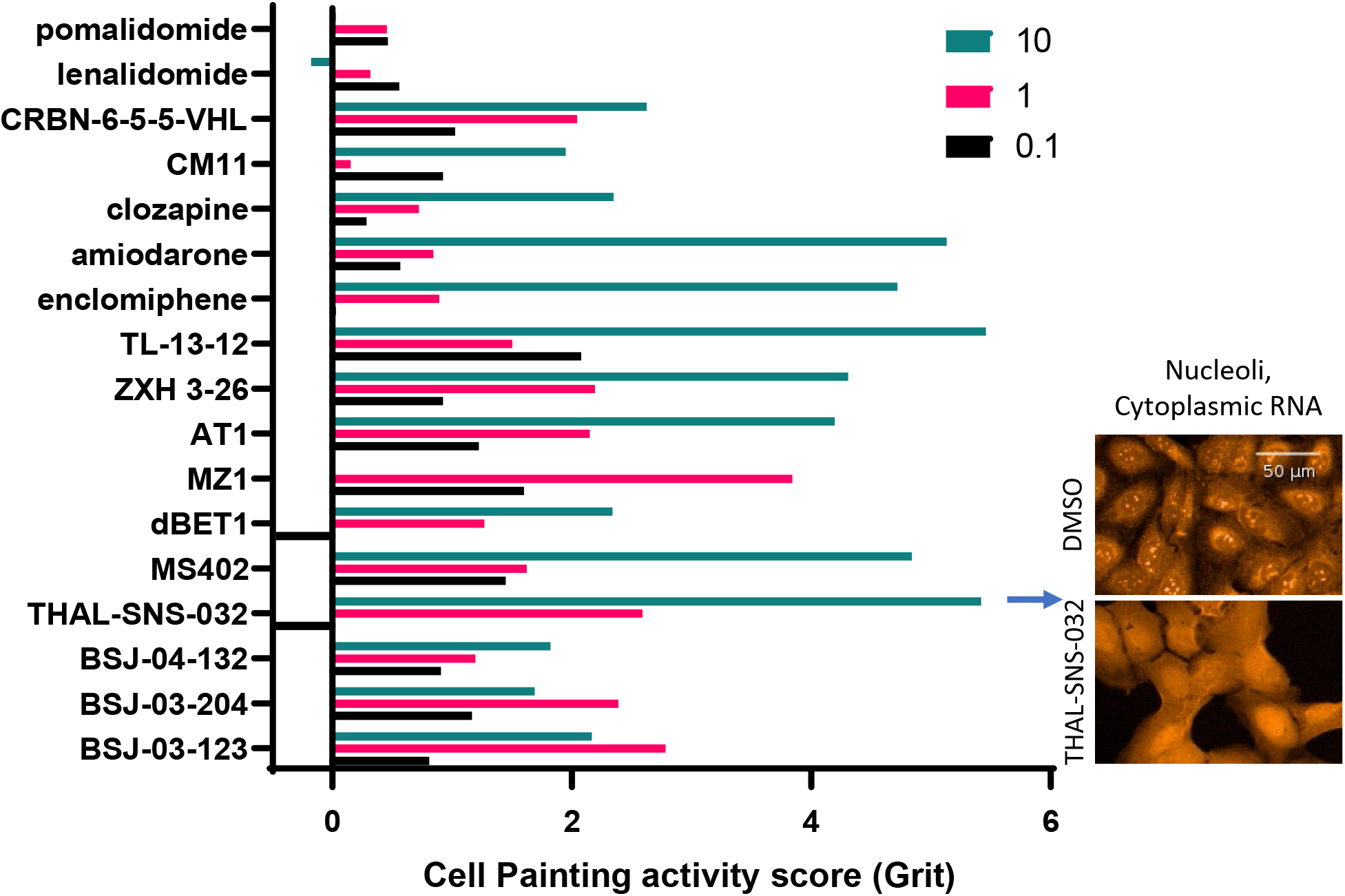
Cell Painting activity score (grit) for published PROTACs and non-PROTACs compounds. The published non-PROTACs compounds dataset consists of commonly used compounds as E3 ligands parts for PROTACs and 3 approved drugs (amiodarone, clozapine and acetaminophen).

The other main observation was that the activity of a PROTAC compound did not always corelate with the activity of the individual PROTAC components. PROTACs are bifunctional molecules containing a binder for the target of interest and a binder for an E3 ligase, the two attached together via a linker. Most of the PROTACs developed at present use the CRBN or VHL E3 ligases. Binders of CRBN include the clinically approved drugs IMiD (immunomodulatory drugs) like lenalidomide and pomalidomide. These two IMiD drugs showed no activity in the Cell Painting assay (Table 1 and Figure 2). However, we did at times observe activity of PROTACs even though the primary target was not expressed in U-2OS cells and no activity was observed with the corresponding E3 binder (warhead) or binder to the target protein (POI, protein of interest) (Figure S2). Hence, this observation illustrated that PROTAC activity can be more than simply the sum of its parts.

### Cell Painting projection revealed different PROTAC signatures

Next, a dimensionality reduction of the PROTACs-Cell Painting profiles was performed with UMAP (Uniform Manifold Approximation and Projection) (McInnes et al., 2018) to understand which phenotypic responses are clustered together using Cell Profiling readouts using this method. The results of this analysis are shown in Figure 3, which suggested a range of different, distinguishable Cell Painting signatures for PROTACs targeting various targets (Figure 3). Furthermore, chemical clustering varied with the concentration of PROTAC used and the Cell Painting activity score (1 vs 10 μM; Figure 3). Looking at specific compounds targeting BRD4, the small molecule inhibitor MS402 clustered together with BRD4 targeting PROTACs, suggesting a similar mode of action (Figure 3, orange annotation). Interestingly, PROTACs from different projects clustered to different regions, suggesting a different mode of action, one in particular clearly showed a different clustering (Figure 3, turquoise -Target 2). The activity of PROTACs targeting Target 2 could not be explained by the E3 ligase or the binder component to the target protein, which is not expressed in U-2 OS cells (Figure 2 and S2), suggesting an off-target mechanism. This observation let us to investigate whether we could link the Cell Painting signature of these PROTACs to a safety finding.

**Figure 3:**
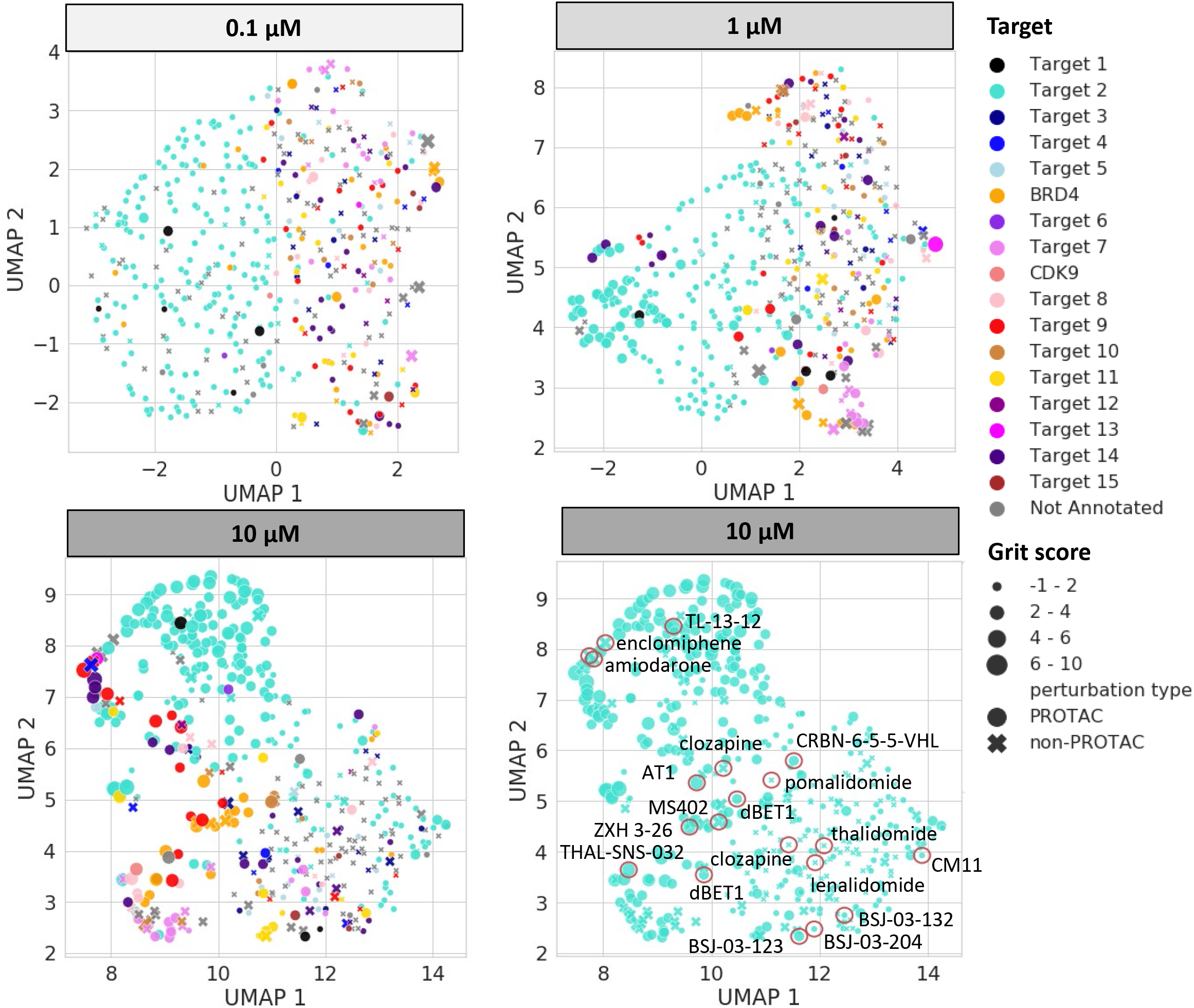
Uniform manifold approximation (UMAP) analysis. UMAP coordinates at concentrations 0.1 1 and 10 μM of all perturbations labelled with the protein that is inhibited or degraded by each non-PROTAC or PROTAC compound respectively. Published PROTACs or non-PROTACs compounds are annotated in the UMAP plot for 10 μM.

### Cell Painting signatures were able to detect activity on mitochondria

To investigate whether Cell Painting profiles could be used to evaluate PROTAC safety liabilities, we employed annotations of *in-vitro* mitotoxicity that were available for part of our compound set. Mitochondrial toxicity annotations for the PROTAC and non-PROTAC compounds were extracted from the Glu/Gal assay (Rana et al., 2018). In this assay, cells are grown in two different media: a high glucose- and galactose-media. Cells grown in high glucose-containing medium use glycolysis for ATP generation and are resistant to mitochondrial insult. Cells grown in galactose-containing medium rely almost exclusively on mitochondria for their ATP production and, hence, are very sensitive to mitochondrial insult (Rana et al., 2018). In total 221 compounds, where 96 were annotated active (mitotoxic) and 125 inactive (not mitotoxic), were used to train the models. Out of the 221 compounds, 149 were PROTACs with 90 having been annotated mitotoxic and 59 having been annotated not mitotoxic. The annotations were further categorised in highly mitotoxic (IC_50_ <1μM; 51 compounds), moderately mitotoxic (IC_50_ between 1 μM and 10 μM; 44 compounds) and not mitotoxic (IC_50_ >10 μM; 126 compounds). At a concentration of 10 μM, the mean grit score was 3.01±1.31, 3.09±1.20, and 1.98±1.59 for highly, moderately, and not mitotoxic PROTACs respectively (Figure 4a). At a concentration of 1μM, the mean grit score was 1.75±0.97, 1.24±0.91 and 1.14±1.28 for highly, moderately, and not mitotoxic PROTACs respectively. The same trend was not observed at concentration 0.1μM, where the mean grit score was 0.64±0.75, 0.73±0.81, and 0.63±0.56 for highly, moderately, and not mitotoxic. Hence, the morphological difference between mitotoxic and non-mitotoxic PROTACs indicated by higher grit scores, is more pronounced at concentrations of 1 and 10 μM. Similar trends were observed for the non-PROTAC compounds (Figure 4a). For example, at concentration 1μM, the mean grit score is 2.36±0.88, 1.36±1.34, and 1.04±1.34 for highly, moderately, and not mitotoxic non-PROTAC compounds respectively. A UMAP dimensionality reduction was performed on the morphological feature space which revealed a separation of mitotoxic compounds from not mitotoxic compounds for both PROTACs and non-PROTACs. Again, this was more evident for the concentration of 10 μM and 1 μM (Figure 4b-d). In addition, we observed a similar signature between the PROTACs active on mitochondria and small molecules that have shown mitochondrial toxicity such as enclomiphene and amiodarone, suggesting a similar mode of action (Figure 3). In summary, our results indicate that mitotoxic compounds induce distinct phenotypic changes which are picked up by the Cell Painting assay, and which might be used to differentiate between mitotoxic and not mitotoxic compounds.

**Figure 4:**
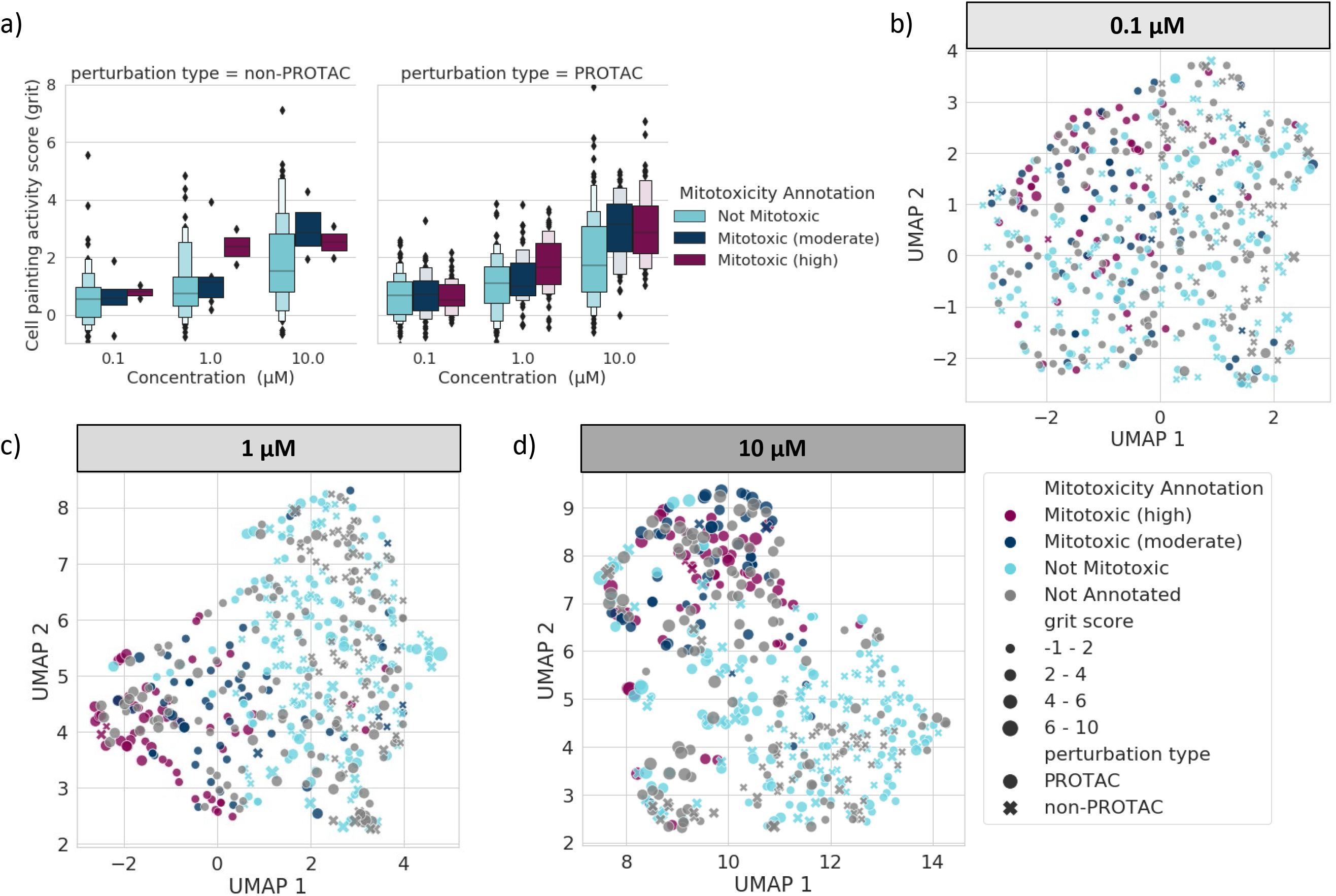
Cell Painting activity with mitochondrial toxicity assay endpoint. a) Cell Painting activity score in the form of grit score across concentrations equal to 0.1, 1.0 and 10.0 μM and labelled based on a mitochondrial toxicity assay endpoint. Uniform manifold approximation (UMAP) coordinates of all perturbations labelled with mitotoxicity annotations at concentrations b) 0.1, c) 1 and d) 10 μM.

### Machine Learning models showed good prediction of mitochondrial toxicity

To investigate whether the Cell Painting profiles can be used as a descriptor for *in-silico* Machine Learning models for mitochondrial toxicity prediction, the profiles were used to train models with three different algorithms namely, Random Forest (RF), Support Vector Classifier (SVC) and eXtreme Gradient Boosting (XGB). Model evaluation results are shown in Figure 5, and the model performance resulted in a ROC-AUC value of 0.93, 0.93, and 0.80 and a F_1_-score of 0.85, 0.87 and 0.74 for concentrations of 10, 1 and 0.1 μM respectively when RF was used (Figure 5a, 5b, 5c). To further validate that the performance is not random, we evaluated whether the models perform better than random models by applying y-scrambling. The y-scrambled models scored a mean ROC-AUC across all algorithms equal to 0.50, 0.51. and 0.49 for concentrations 0.1, 1 and 10 μM respectively (i.e., close to the expected value of 0.5) as shown in Figure S3. Hence, the models perform significantly better than the y-scrambling models and thus they are unlikely to have been obtained by chance.

**Figure 5:**
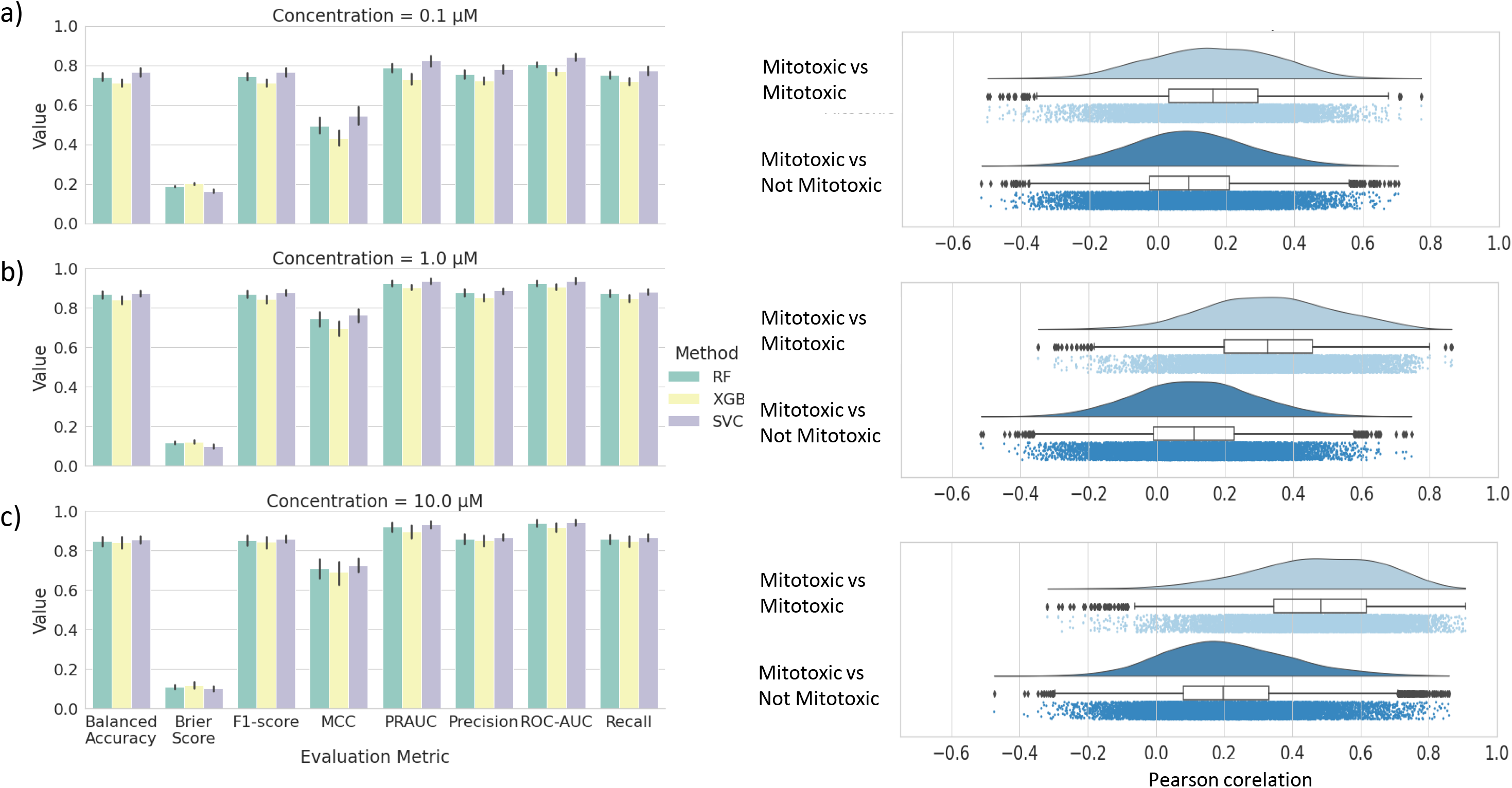
Performance of models for mitochondrial toxicity prediction. Mitochondrial toxicity prediction performance using the Cell Painting features and three different algorithms; RF, XGB and SVC at concentrations a) 10, b) 1 and c) 0.1 μM. The error bars correspond to the confidence interval across all splits and random states used for cross validation. Intra-class (Mitotoxic vs Mitotoxic) vs Inter-class (Mitotoxic vs not Mitotoxic) Pearson correlation of the image-based features are shown for each concentration.

Using the SVC algorithm, the balanced accuracy was equal to 0.76, 0.88 and 0.85 when the models were trained with profiles from concentrations 0.1, 1 and 10 μM respectively (Figure 5a, 5b, 5c). Hence, models trained with Cell Painting profiles from the two higher concentrations of 1 and 10 μM outperformed the models trained on profiles from the concentration of 0.1 μM. Similarly, concentrations of 1 and 10 μM outperformed the concentration of 0.1 μM regardless of the algorithm used as shown in Figure 5. This is in agreement with the finding described above that grit scores were larger for mitotoxic compounds at the two higher concentrations than at the lower concentration tested. Furthermore, this can be explained by the fact that, a high intra-class correlation was observed between the mitotoxic compounds in the Cell Painting features at a concentration of 10 and 1 μM with a median of 0.48 and 0.32 respectively, compared to a lower intra-class Pearson correlation at concentration of 0.1 μM with a median of 0.16 (Figure 5a, 5b, 5c). Hence, PROTACs and compounds that cause mitochondrial toxicity are significantly more similar to each other at concentrations 1 and 10 μM (Figure 5b, 5c), compared to features derived at 0.1 μM (Figure 5a). Furthermore, a high difference in the intraclass and interclass correlations (between mitotoxic and not mitotoxic) were observed and were equal to 0.28, 0.21 and 0.07 for concentration 10, 1 and 0.1 μM respectively. Overall, this means that active compounds at concentrations of 10 and 1 μM are clearly different from inactive compounds (median similarities of 0.48 vs 0.20 and 0.32 vs 0.11 respectively), while being less indistinguishable at concentration 0.1 μM (median similarities of 0.16 vs 0.09). Taken together, this similarity analysis additionally explains why using concentrations 1 and 10 μM outperforms concentration 0.1 μM model performance.

### Prospective experimental model validation

To further validate our findings, we performed external validation for our mitochondrial toxicity models. Out of the total PROTACs and compounds tested with in the Glu/Gal assay, there were 39 PROTACs that were tested later out of which five were mitotoxic and 34 were not mitotoxic, which were used as a prospective test set. A similarity analysis (by calculating Pearson correlation) was initially performed between the 39 query PROTACs to the compounds which cause mitochondrial toxicity and those which do not (i.e., the compounds in the models).For concentrations 1 and 10 μM, the mitotoxic query PROTACs show a higher correlation with the mitotoxic compared to the correlation with the not mitotoxic (Figure S4). In addition, the not mitotoxic query PROTACs do not show a high correlation with the mitotoxic PROTACs in the models (Figure S4). This supported our assumption that the models would be able to also classify the prospective test set correctly.

The mitochondrial toxicity of the 39 PROTACs was hence predicted by all the models and the external validation results are summarised in Figure 6. In addition, results are summarised with confusion matrices and model evaluation metrics in Figures S5 and S6 respectively. The models trained with data at concentration 1 and 10μM performed well and outperformed the models trained with data at a concentration of 0.1 μM. For example, the balanced accuracy was equal to 0.68, 0.96 and 0.89 when the models were trained with profiles from concentrations 0.1, 1 and 10 μM respectively (Figure S6). Moreover, the models trained with the data at a concentration of 0.1μM showed relatively high retrieval for mitotoxic PROTACs (more than 60% of mitotoxic PROTACs were correctly classified, (Figure 6a), but on the other hand showed high false-positive rates (Figure S5). The models trained with the data from concentration 1 and 10μM were consistently able to predict the majority of the mitotoxic PROTACs (Figure 6a), with the models using data from the concentration of 1 μM being able to predict 100% of the mitotoxic PROTACs, regardless of the algorithm used. Models trained with the data from the highest concentration of 10 μM are able to correctly detect 60%, 80% and 80% of the mitotoxic PROTACs using the RF, SVC and XGBOOST algorithms respectively (Figure 6a). On the other hand, the models trained with data from concentration 10 μM have a lower number of false positives and thus a higher number of true negatives compared to models trained with data from concentration 1 μM (Figure S5). Finally, 97% and 91-97% of the not mitotoxic PROTACs are correctly classified using the models trained with data from concentration 10 and 1 μM respectively (Figures 6b and S5).

**Figure 6:**
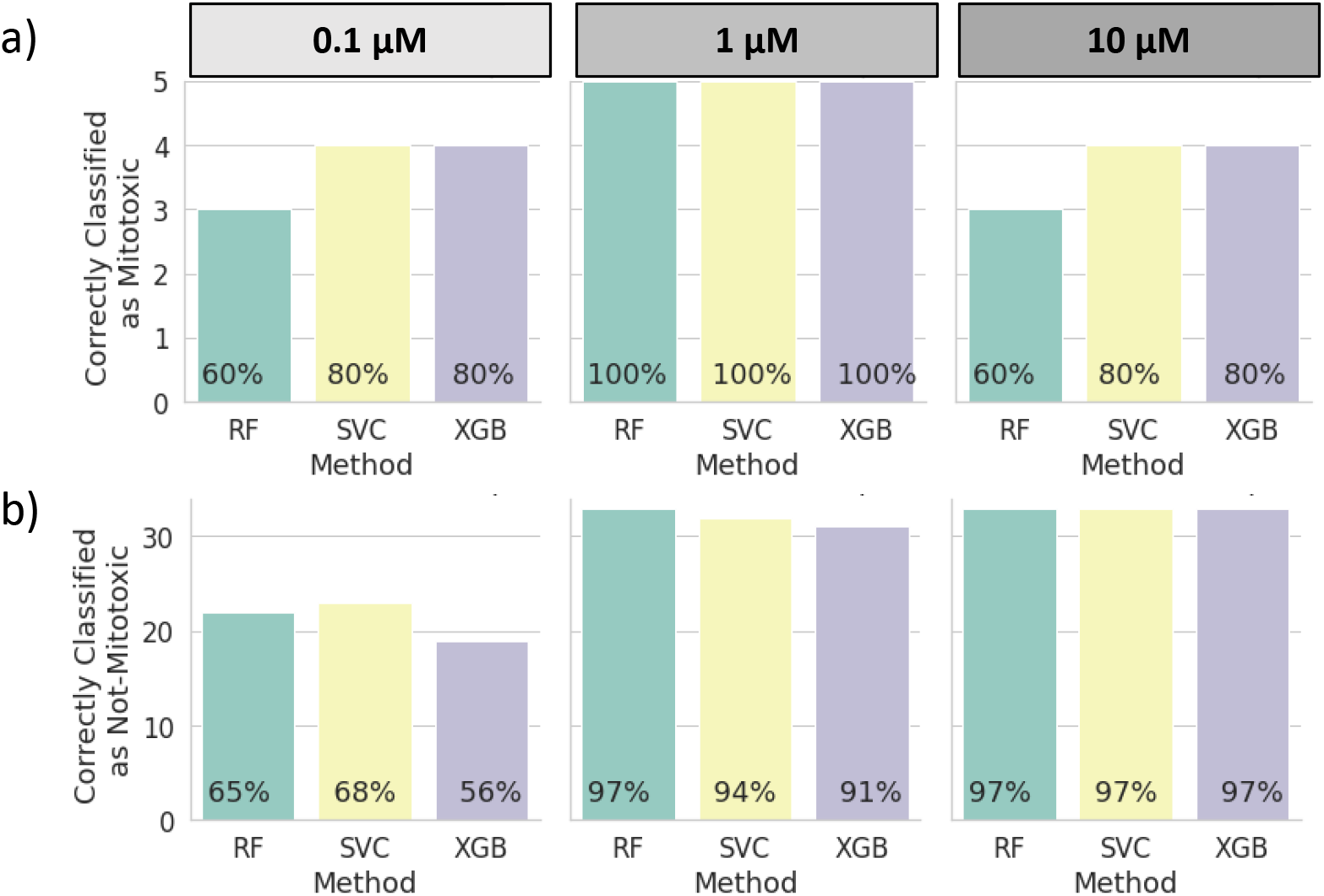
Prospective experimental model validation. Number (and percentage) of correctly classified a) mitotoxic and b) not-mitotoxic PROTACs, obtained with the models trained with RF, SVC and XGB algorithms and with data from concentration 0.1, 1 and 10 μM.

## Discussion

The increasing interest in PROTACs as a novel therapeutic modality, results in the need for assays to profile these bRo5 compounds. Therefore, in this work, the Cell Painting assay was used to profile a series of PROTAC and non-PROTAC compounds from various projects based on the hypothesis that Cell Painting assay could quantitatively study the morphological impact of PROTACs. Two different metrics, a Euclidean distance–based and the grit score, revealed that PROTACs’ and non-PROTACs’ profiles are different from the neutral controls and thus the Cell Painting assay was able to capture morphological changes induced by PROTACs. In addition, Euclidean distance-based method and grit score revealed a higher number of active compounds on the Cell Painting assay and a stronger phenotypic effect respectively as the concentration of compounds was increasing.

Focusing on particular examples from published PROTACs, we found that PROTACs degrading targets such as BRD4 show an activity on the Cell Painting assay. In addition, a PROTAC targeting CDK9 (THAL-SNS-032) showed a high activity and by looking at the raw images, the phenotype that was observed was consistent with the function of CDK9 in cell cycle progression. More surprisingly, we observed that the activity of a PROTAC on the Cell Painting assay did not necessarily correlate with the activity of its individual components (i.e., POI ligand and E3 ligase ligand). This observation highlighted that PROTACs’ activity on Cell Painting assay is not just the sum of its parts. Furthermore, upon a dimensionality reduction of the PROTACs-Cell Painting profiles with UMAP, we were able to understand whether and which phenotypic responses are clustered together given the target they degrade. Results suggested a range of different and distinguishable Cell Painting signatures for PROTACs targeting various targets such as the BRD4. Looking at specific compounds targeting BRD4, the small molecule inhibitor MS402 clustered together with BRD4 targeting PROTACs, suggesting a similar mode of action.

However, there were cases, where PROTACs showed a Cell Painting activity even though the primary target was not expressed in U2OS cells and no activity was observed with the corresponding binder to the target protein. This was an indication that this effect could be related to a PROTACs’ off-target effect and thus could be a useful information to better understand PROTACs’ safety profiles. Therefore, we trained *in-silico* machine learning models to predict compounds’ (including PROTACs) mitochondrial toxicity using the Cell Painting profiles as descriptors for Random Forest, Support Vector Classifier and XGB algorithms. Models trained with the Cell Painting features at concentration of 1 and 10μM outperformed the performance of 0.1 μM. In addition, a models’ prospective validation was performed showing that the models trained with data at concentration 1 and 10 μM performed well. Mitochondrial toxicity is a major safety concern associated with serious organ toxicities and a frequent cause of late-stage drug withdrawals. With the growing presence of new modalities, including PROTACs, there is an urging need to evaluate such safety risks for novel compounds. Numerous efforts exist to evaluate or predict small molecule’s mitochondrial toxicity and different assays have been developed capturing various mechanisms of drug-induced mitochondrial toxicity including the Glu/Gal assay used here (Will and Dykens, 2014). However, Hynes et al., 2013, showed that the Glu/Gal assay only detects about 2 - 5% of all mitotoxicants, which further highlights the reality that most compounds that cause organ toxicity do so via multiple off-target mechanisms. Our study highlighted the potential to use Cell Painting for mitotoxicity prediction and given its throughput could be used a very useful method to screen compounds at scale, including new modalities such as PROTACs.

## Significance

In this work, we evaluated whether PROTACs can be profiled with the Cell Painting assay. In addition, it was evaluated whether the cell morphological profiles derived from the Cell Painting assay could be used as a PROTACs’ descriptor to predict mitochondrial toxicity. Results showed that PROTACs can induce cell morphological changes, and this was proved by using two different metrics: Euclidean distance-based and grit score. In addition, the PROTACs – Cell Painting profiles were used as descriptors in mitochondrial toxicity prediction models and resulted in models with high performance. Finally, the models showed a good performance in predicting the activity of a prospective validation set of PROTACs. According to our knowledge, this is the first time to show that PROTACs can change the cellular morphology using the Cell Painting assay and this finding creates a new hypothesis on how the readouts from this assay can be used to better understand this new data modality.

## Methods

### Cell Culture and Seeding

U-2 OS cells, a human osteosarcoma cell line, were sourced from AstraZeneca’s Global Cell Bank (ATCC Cat# HTB-96). Cells were cultured in McCoy’s 5A media (Gibco, #26600023) supplemented with 10% (v/v) foetal bovine serum (Gibco, #10270106) at 37°C, 5% (v/v) CO2, 95% humidity. After reaching *ca*. 80% confluency, cells were washed with PBS (Gibco, #10010056) then detached from culture flasks using TrypLE Express enzyme (Gibco, #12604013) and resuspended in McCoy’s media. Cells were counted using a Vi-CELL (Beckman Coulter, #383556) then diluted with McCoy’s media to achieve a count of 1,250 cells per well using a dispense volume of 40 μL per well. The cell suspension was dispensed into CellCarrier-384 Ultra microplates (Perkin Elmer, #6057300) using a Multidrop™ Combi (ThermoFisher, #5840300) with a standard-tube cassette (ThermoFisher, #24072670). Microplates were left at room temperature for 1h before transferring to a SteriStore (HighRes Biosolutions) microplate incubator at 37°C, 5% (v/v) CO2, 95% humidity for 24h prior to compound addition.

### Compound Treatment

PROTACs were sourced internally through the AstraZeneca Compound Management Group. PROTACs were prepared as 10, 1 and 0.1 mM source stocks respectively (in DMSO) and plated into intermediate 384-well echo-qualified source plates (Labcyte, #PP-0200). 24h post-seeding, assay plates were dosed using an Echo® 655T acoustic dispenser (Labcyte) from the appropriate compound stock to perform a 1000-fold dilution, to achieve assay concentrations of 10, 1 and 0.1 μM respectively. Where required, assay wells had DMSO added to maintain a final DMSO concentration of 0.1% (v/v). Assay plates were returned to the SteriStore incubator for a further 48h prior to performing the cell staining protocol.

### Cell Staining

The Cell Painting staining procedure was performed according to the protocol by Bray et al., 2016 with some adjustments to stain concentrations and methodology. Hanks’ balanced salt solution (HBSS) 10x was sourced from AstraZeneca’s media preparation department and diluted in dH_2_O then filtered using a 0.22μm filter (Corning, CLS430517). MitoTracker stain (ThermoFisher, M22426), was prepared as a 1mM stock solution in DMSO and then made up as a working stain solution in McCoy’s 5A medium, at a final concentration of 0.5 μM. The remaining stains were prepared in 1% (w/v) bovine serum albumin (BSA) (Sigma Aldrich, A4503) in 1x HBSS containing 0.1% (v/v) Triton X-100 (Sigma Aldrich, T8787).

Following compound incubation, 10μl of MitoTracker working solution was added to the plate and incubated for 30 minutes at 37°C, 5% CO_2_, 95% humidity. The following steps were all carried out at room temperature in the dark. Cells were fixed by adding 25 μl of 12% v/v formaldehyde in PBS (to achieve final concentration of 3.25% v/v). Plates were incubated for 20 min then washed using a Blue®Washer centrifugal plate washer (BlueCat Bio, Neudrossenfeld, Germany). Following this, 15 μl of stain solution containing 5 μg/ml Hoechst 33342 (ThermoFisher, H3570), 1.5 μg/ml Wheat-germ Agglutinin Alexa Fluor® 555 conjugate (ThermoFisher, W32464), 10 μg/mL ConcanavalinA Alexa Fluor® 488 conjugate (ThermoFisher, C11252), 5μl/mL Phalloidin Alexa Fluor® 568 conjugate (ThermoFisher, A12380) and 9 μM SYTO14 (ThermoFisher, S7576) was dispensed to each well and incubated for 30min then removed prior to a final wash and subsequent addition of 1x HBSS to each well. Plates were sealed and then imaged.

### Imaging

Cells were imaged with a CellVoyager CV8000 (Yokogawa, Tokyo, Japan) using a 20x water-immersion objective lens (Olympus, Tokyo, Japan; NA 1.0). Five imaging channels were acquired to visualise all fluorescent stains: DNA (ex: 405nm, em: 445/45nm), ER (ex: 488nm, em: 525/50nm), RNA (ex: 488nm, em: 600/37nm), AGP (ex: 561nm, em: 600/37nm) and Mito (ex: 640nm, em: 676/29nm). Four fields of view were acquired per well to capture sufficient numbers of cells per perturbation.

### Image Analysis and Feature Extraction

Images were saved as 16-bit .tif files without binning (1994 x 1994 pixels). Images were analysed using CellProfiler™ biological image analysis software (v 4.0.7). The segmentation of individual nuclei was performed using the DNA channel and subsequent cellular segmentation using the AGP channel. Cells touching the boundary of the image were excluded from subsequent analysis. A total of 4700 features were calculated, relating to either whole-image level properties or individual objects (cells, nuclei or cytoplasm). Features include pixel intensity co-localisation measurements; granularity and textural measurements of objects taken across a range of pixel distances; the presence and proximity of neighbouring objects; the distribution of staining intensity patterns and size/shape metrics.

### Data Curation and Normalisation

A normalisation process was applied as described by Way et al., 2020, since HTS experiments can be affected by systematic row, column and edge effects (Makarenkov et al., 2007) and thus, there is a need for data normalisation in order to reduce false positives in such experiments (Dragiev et al., 2011, 2012). Firstly, single cell data per well were merged by calculating their median value. Next, data were normalised using the median and the median absolute deviation (MAD) of feature values from empty wells (DMSO) as the centre and scale parameters respectively. We normalised all perturbation profiles by subtracting the centre (median) and dividing by the scale (MAD) and did for each plate individually.

### Feature Selection

A feature selection was performed to remove features based on a set of criteria. The first criterion was the variance of the features across profiles and hence features with a variance less than 1 were removed. In addition, features with a high standard deviation were filtered out and we used a standard deviation threshold equal to 20. According to Way et al., 2020, features with a high standard deviation after normalisation are considered as feature outliers and should be removed. In addition, features with missing values in any profile were filtered out. Moreover, pairwise correlations were calculated for all the features and randomly removed 1 feature from each pair with a Pearson correlation greater than or equal to 0.9. As a result of these processes, 669 features remained.

### Evaluation of PROTACs activity on Cell Painting assay

Two different methodologies were used to evaluate whether PROTACs were active on the Cell Painting assay screen. The first one was a Euclidean distance-based approach and the second is the calculation of grit score. The first approach was described by Cox et al., 2020, and we used it in order to calculate which PROTACs were “active” on the assay using a 95th percentile cut-off on the null distribution of Euclidean distances between individual DMSO control profiles and the mean DMSO control profile.

In addition, we used the grit score (https://github.com/cytomining/cytominer-eval, https://github.com/broadinstitute/grit-benchmark), which captures the phenotypic strength of a perturbation in a profiling experiment and combines two concepts. The first is the replicate reproducibility and the second is difference from the DMSO control. Firstly, for each target profile (i.e., PROTACs) pairwise Pearson correlations were calculated for both PROTACs replicates and control replicates. Hence, the pairwise correlations form two distinct distributions (replicate and control). Then using the control profiles only, a Z-score transform is obtained, which is then used to transform the PROTACs’ replicates. The mean of PROTACs’ replicates Z-scores are calculated, and this is the final score termed grit score. Since grit is based on Z-scores, the magnitude can be easily compared between perturbations and is a directly interpretable value. For example, a grit score of 3 for a PROTAC X compared to a neutral control means that on average PROTAC X is 3 standard deviations more similar to replicates than to DMSO controls. Therefore, it is considered as the PROTACs’ average reproducibility with respect to the neutral control similarity. Grit score was calculated with the cytominer-eval Python package(https://github.com/cytomining/cytominer-eval), developed by the Broad Institute.

### Glu/Gal Assay for mitochondrial toxicity assessment

This assay is used to assess potential test substances that can trigger mitochondrial dysfunction. HepG2 cells are cultured in a) glucose containing and b) galactose containing media and are exposed for 24 h to a concentration of x of the test compounds. Following treatment, the IC_50_ (μM) galactose is measured, and it corresponds to the average galactose signal value which is halfway between the baseline and the average maximal signal for the substance tested. If IC_50_ (μM) galactose is more than 10 then the substance is considered inactive (i.e., does not cause mitochondrial toxicity) and if less than or equal to 10, then it is active and causes mitochondrial toxicity. This mitochondrial toxicity annotation was used to train predictive models for PROTACS’ mitochondrial toxicity prediction. In total 221 compounds (PROTAC and non-PROTAC) were used to train the models with 96 active (mitotoxic) compounds and 125 inactive (not mitotoxic) compounds. Out of the total of 221 compounds, the 149 were PROTACs and in more detail, the 90 PROTACs were mitotoxic with the rest of PROTACs being not mitotoxic.

### Mitochondrial toxicity *in-silico* model training and evaluation

Three times nested five-fold cross-validation was performed with the StratifiedShuffleSplit Python function from Scikit-Learn (Pedregosa et al., 2011). The Stratified Shuffle Split (SSS) splits a dataset into a train and test set by preserving the same percentage of data for each class (active and inactive) as in the initial dataset. Schematic representation of model training process is shown in Figure S7.

Initial data were splitted in 70% train and 30% test set respectively 5 times using the stratified shuffle split function from Scikit-Learn. The training set was furhter splitted 5 times using the stratified shuffle split function from Scikit-Learn to identify the optimal hyperparameters using hyperopt and cross validation score function from Scikit-Learn. When hyperparameters were selected, the models were trained and the compounds in the test set were predicted. This process was repeated with 3 different random seeds when the initial data was splitted.

Machine Learning models to predict PROTACS’ mitochondrial toxicity were trained with three different algorithms: a) Random Forest (RF), Support Vector Classifier (SVC) and c) XGBOOST (Chen and Guestrin, 2016). RF and SVC were implemented with the RandomForestClassifier and SupportVectorClassifier functions respectively from Scikit-Learn (Pedregosa et al., 2011) and eXtreme Gradient Boosting (XGB) with the XGBClassifier from xgboost python package. Hyperparameter selection for each of the algorithms was performed by using hyperopt python package(Bergstra et al., 2013, 2015). The parameters and the range of values (configuration space), which were explored for each algorithm are included in Supplementary Information (Table S1). Cell Painting features were used as descriptors for the models. We used model evaluation metrics from Scikit-Learn, which were averaged to give the overall performance across the different folds of cross-validation for Receiver Operating Characteristic - Area Under Curve (ROC-AUC), Precision (Equation 1), Recall (Equation 2), F_1_-score (Equation 3), Balanced accuracy (Equation 4), brier score (Equation 5) and Mathews Correlation Coefficient (MCC, Equation 6).

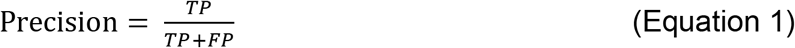

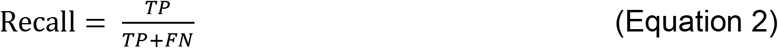

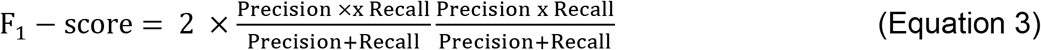

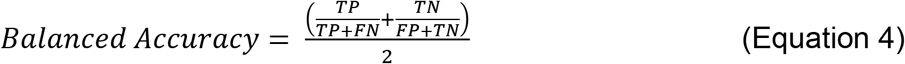

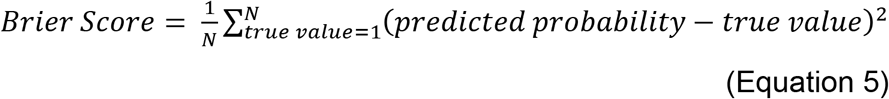

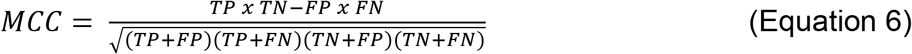

TP denotes true-positives, FP denotes false-positives, TN denotes true-negatives and FN denotes false-negatives.

Finally, y-scrambling (Lipiński and Szurmak, 2017) was performed in order to evaluate whether the trained models performed better than the y-scrambled models. Y-scrambling was applied by randomly reorganising the mitochondrial toxicity labels. Models were rebuilt and evaluated with the same parameters as the unscrambled (actual) models.

### Prospective Model Validation

PROTACs that have been tested on the mitochondrial toxicity assay after the PROTACs, which have been included in the benchmarking of models were extracted and used as prospective validation set. This set included 5 PROTACs that caused mitochondrial toxicity and 34, which did not.

## Acknowledgments

We thank our AstraZeneca colleagues for the internal reviewing and the helpful discussion during the preparation of the manuscript.

## Author contributions

Conceptualization, E.Mo., E.Mu., K.M.; Methodology, M.-A.T., E.Mo, G.W., T.M, R.T., F.M., A.M., L.M.; Biological Research E.Mo, G.W, K.J, T.M; Data analysis, G.W, K.M., M.-A.T.; Writing, K.M, E.Mo, M.-A.T.; Supervision, K.M., A.B., I.B., O.E., L.M. All authors discussed the results and commented on the manuscript.

## Declaration of interest

All the authors are employees of AstraZeneca.

## Supplemental information

**Figure S1:**
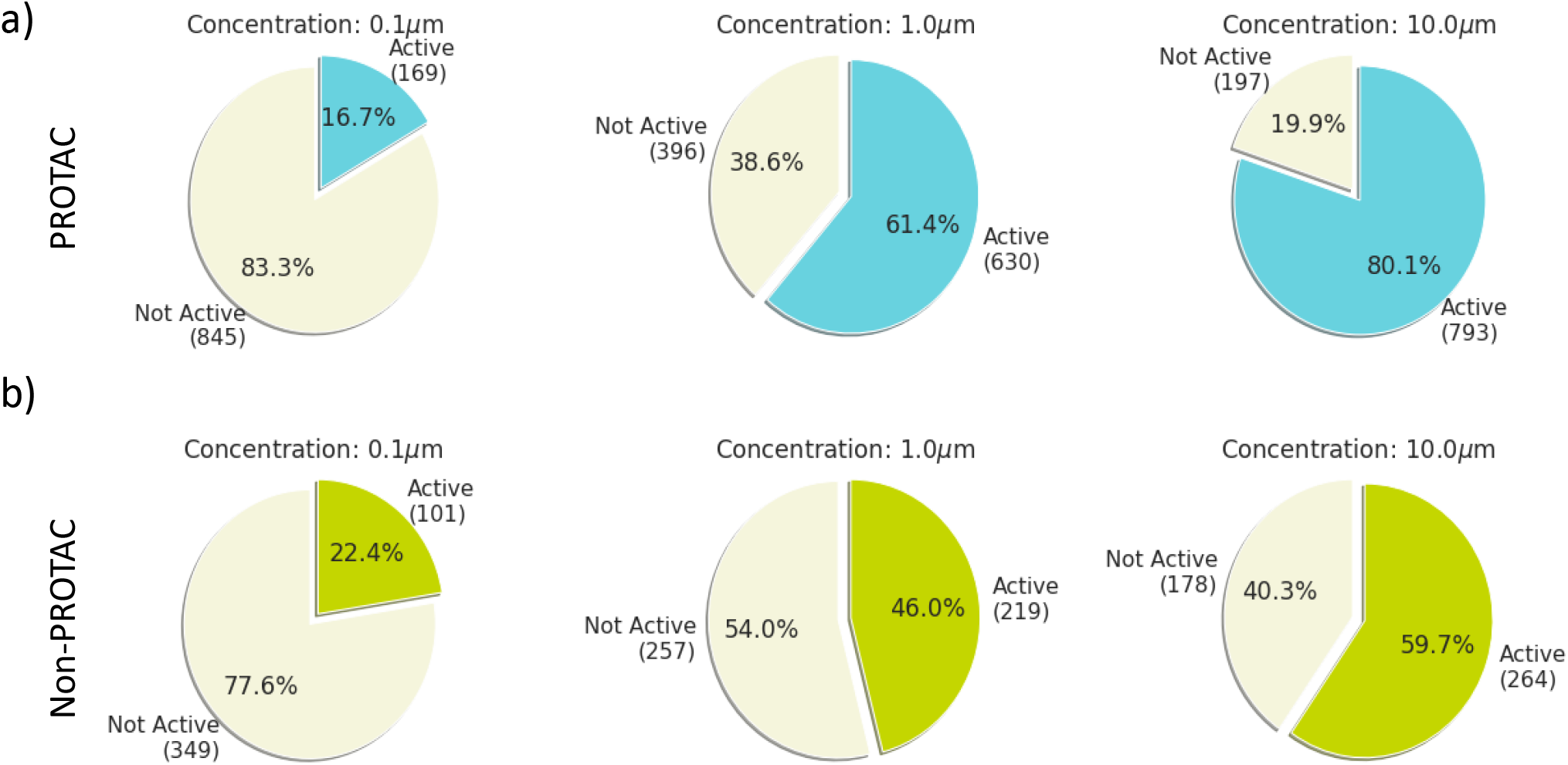
Cell Painting score with the Euclidean-based method. Percentage of a) PROTACs and b) non-PROTACs compounds identified as active on the Cell Painting assay with the Euclidean-based method (i.e. compounds that are able to change the cellular morphology) at concentration of 0.1, 1 and 10 μM. Euclidean distance-based method showed that the number of active compounds increases as the concentration increases.

**Figure S2:**
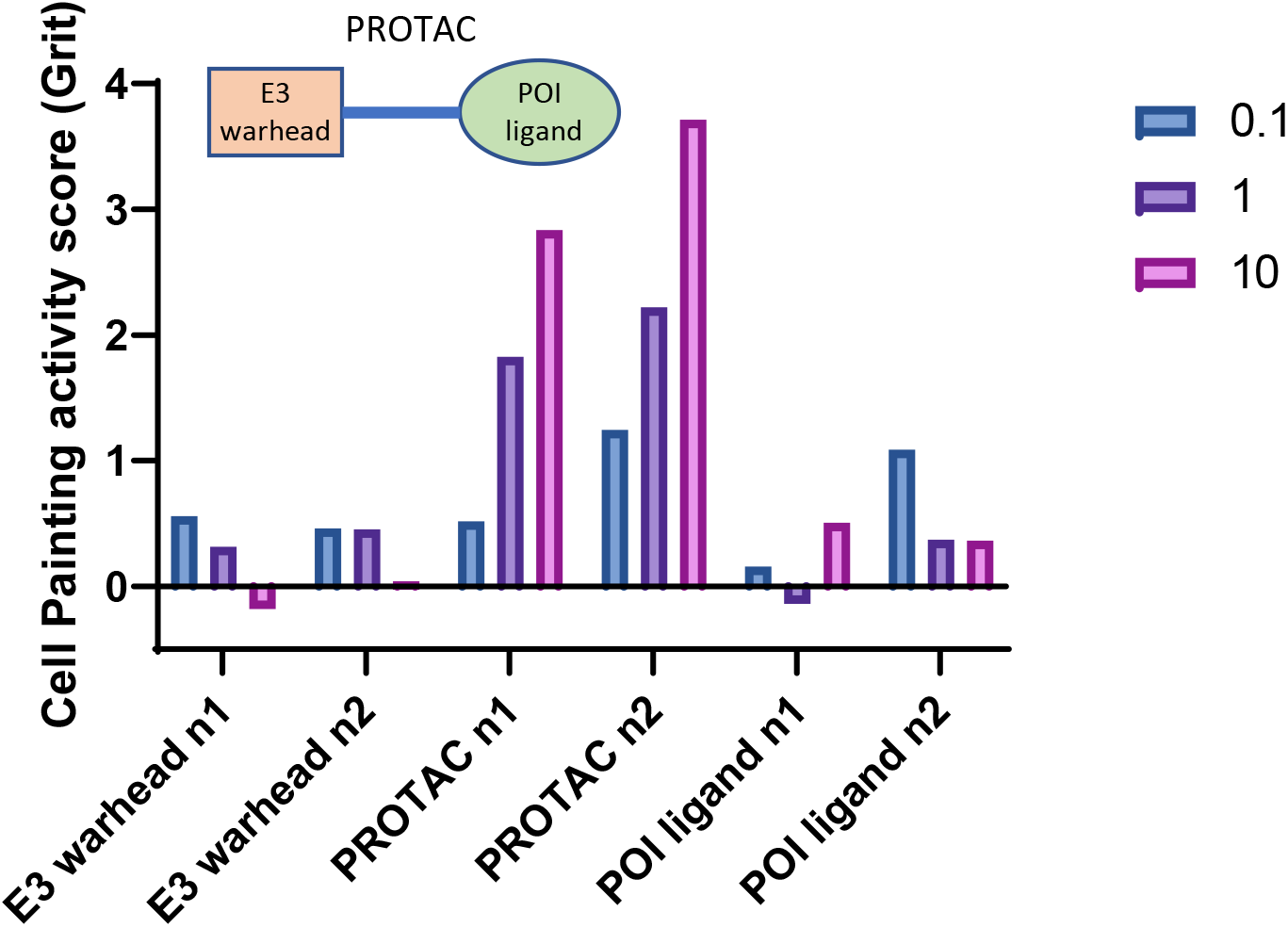
Cell Painting activity for individual PROTAC components. Cell Painting activity score (Grit) for two PROTACs together with their corresponding part (E3 warheads and protein of interest POI ligands).

**Figure S3:**
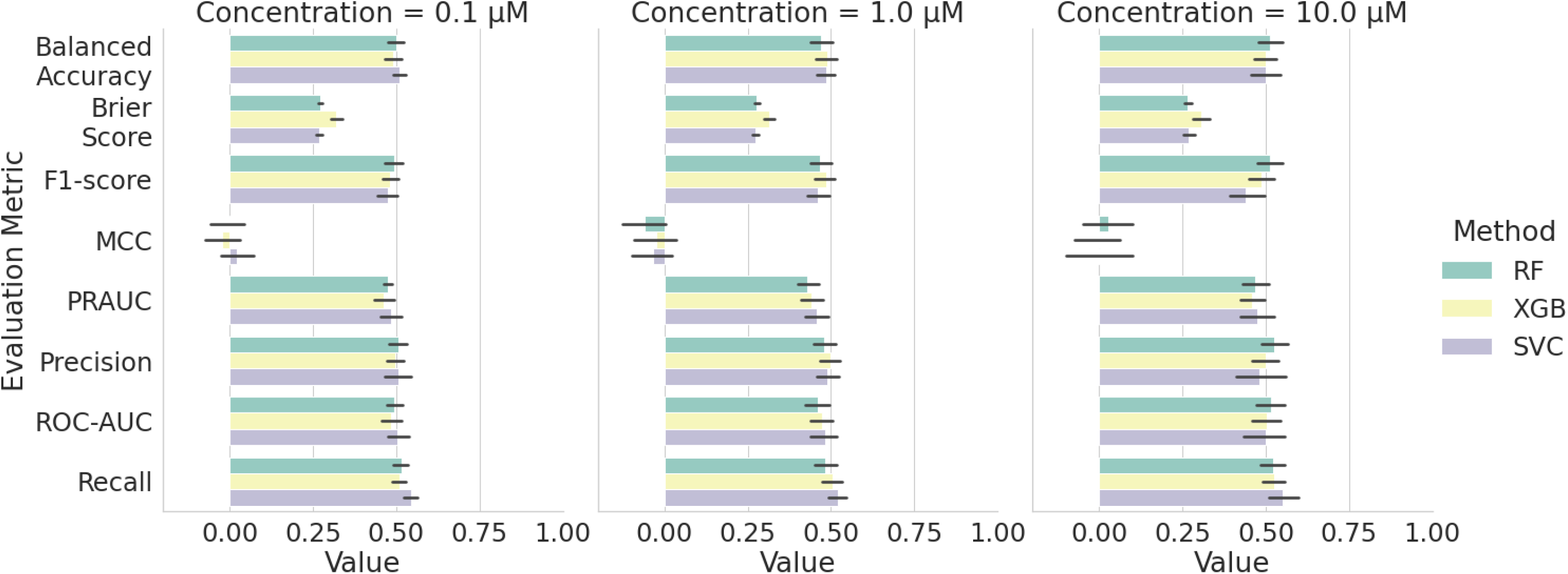
Performance of y-scrambled models. Performance of y-scrambled models for mitochondrial toxicity prediction using the Cell Painting features and three different algorithms; RF, XGB and SVC at concentrations 0.1, 1 and 10 μM. The error bars correspond to the confidence interval across all splits and random states used for cross validation.

**Figure S4:**
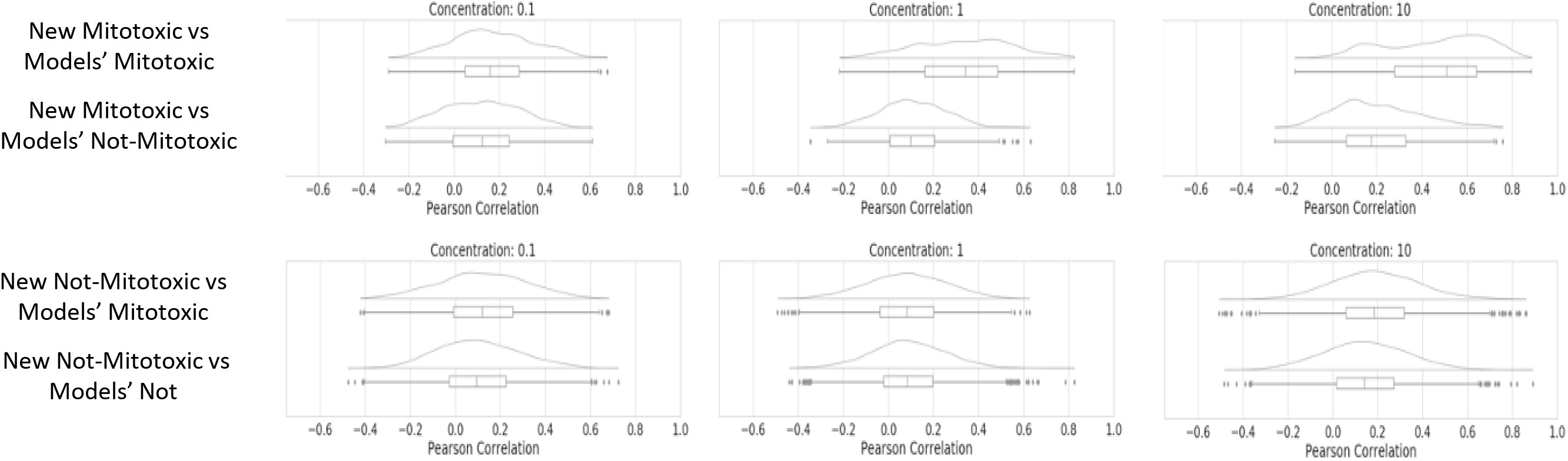
Pairwise Pearson correlation. Pairwise Pearson correlation in the Cell Painting features space between the PROTACs in the external validation set and the compounds (PROTACs and non-PROTACs) in the mitochondrial toxicity models. The four following comparisons are performed. “New Mitotoxic vs Models’ Mitotoxic” corresponds to the pairwise Pearson correlation calculation between the mitotoxic PROTACs in the external validation set and the mitotoxic compounds in the model. “New Mitotoxic vs Models’ Not-Mitotoxic” corresponds to the pairwise Pearson correlation calculation between the mitotoxic PROTACs in the external validation set and the not-mitotoxic compounds in the model. “New Not-Mitotoxic vs Models’ Mitotoxic” corresponds to the pairwise Pearson correlation calculation between the not mitotoxic PROTACs in the external validation set and the mitotoxic compounds in the model. “New Not-Mitotoxic vs Models’ Not-Mitotoxic” corresponds to the pairwise Pearson correlation calculation between the not-mitotoxic PROTACs in the external validation set and the not-mitotoxic compounds in the model. These calculations are performed for concentration a) 0.1, b) 1 and c) 10 μM.

**Figure S5:**
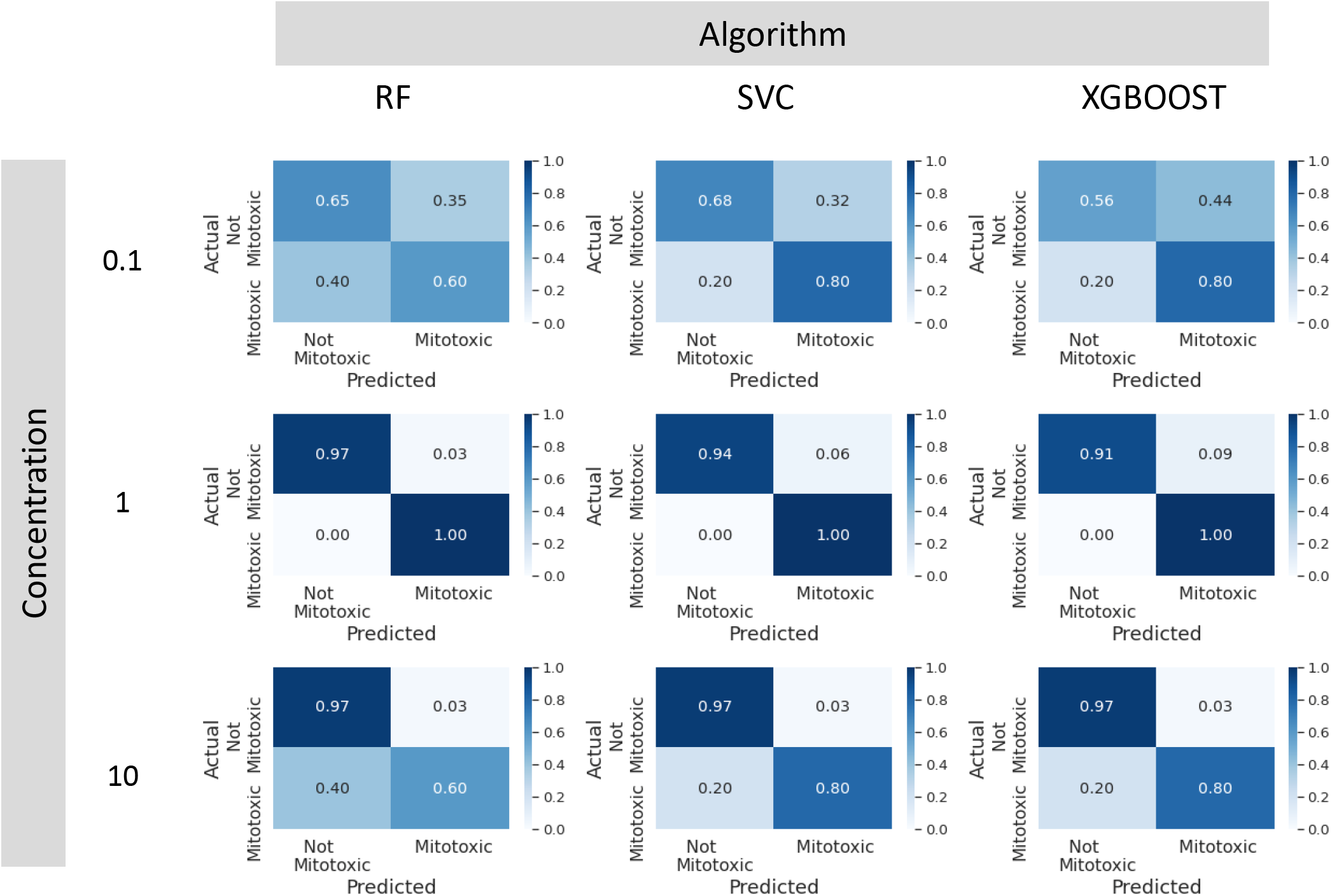
Prospective experimental model validation results visualised with confusion matrices. Results obtained with the models trained with RF, SVC and XGB algorithms and with data from concentration 0.1,1 and 10 mM.

**Figure S6:**
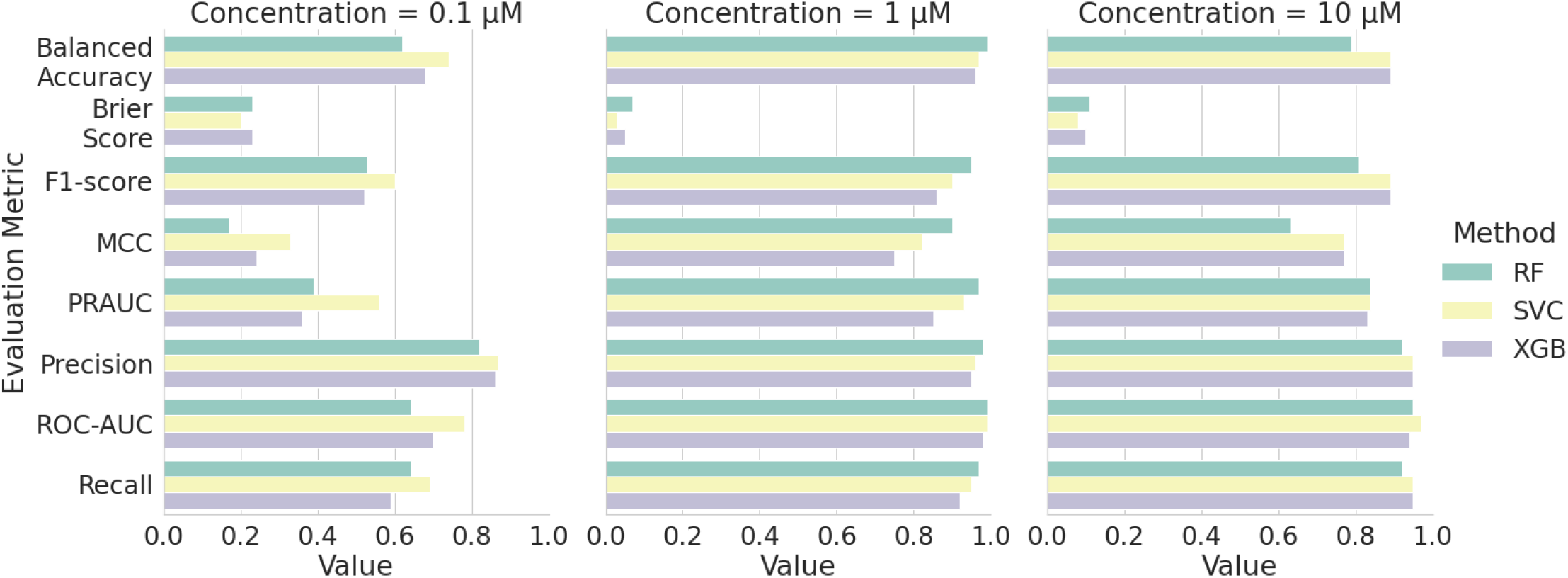
Model performance on the prospective validation set assessed with a range of evaluation metrics. Mitochondrial toxicity prediction performance using the Cell Painting features and three different algorithms; RF, XGB and SVC at concentrations a) 10, b) 1 and c) 0.1 μM. The error bars correspond to the confidence interval across all splits and random states used for cross validation.

**Figure S7:**
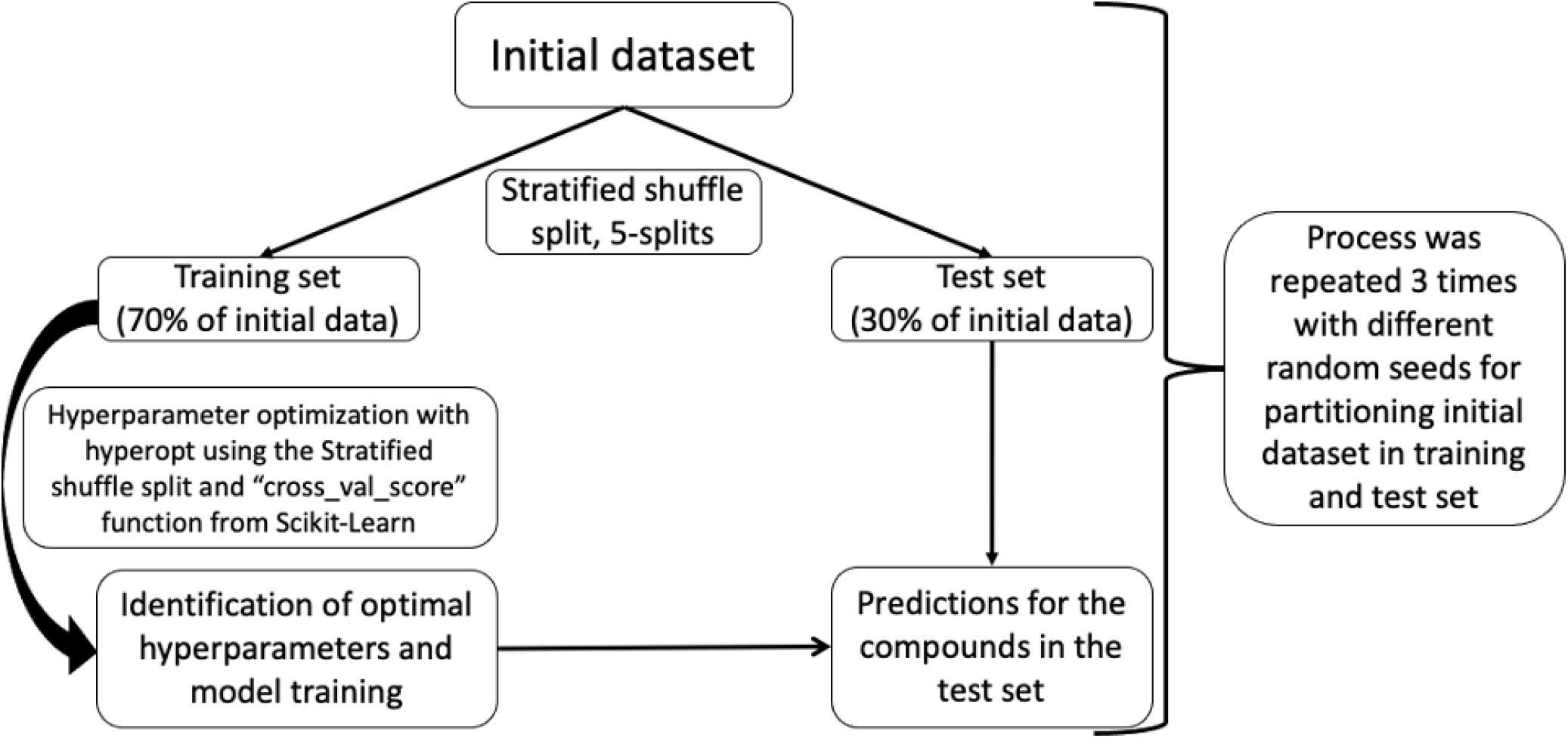
Schematic representation of model training process. Initial data were partitioned in 70% train and 30% test set respectively, 5 times using the stratified shuffle split function from Scikit-Learn. The training set was further partitioned 5 times using the stratified shuffle split function from Scikit-Learn to identify the optimal hyperparameters using hyperopt and cross validation score function from Scikit-Learn. When hyperparameters were selected the models were trained and the compounds in the test set were predicted. This process was repeated with 3 different random seeds when the initial data were partitioned.

**Table S1:** Considered machine learning hyperparameters. Range of hyperparameters’ values considered for the RF, SVC and XGB algorithms. Hyperparameters were systematically evaluated using hyperopt python package.

| Algorithm | Hyperparameter | Values |
| --- | --- | --- |
| Random Forest | max_depth | 3-50 with increments of 1 |
|  | n_estimators | 100-1000 with increments of 100 |
|  | min_samples_split | 2-50 |
|  | min_samples_leaf | 1-15 with increments of 1 |
| Support Vector Classifier<br>(with rbf kernel) | gamma | $10^{-10}$-1 |
| | C | $10^{-4}$-1000 |
| eXtreme Gradient Boosting (XGB) | max_depth | 3-18 with increments of 1 |
|  | n_estimators | 100-1000 with increments of 100 |
|  | gamma | 0-9 |
|  | reg_alpha | 0.1-100 with increments of 1 |
|  | colsample_bytree | 0-1 |
|  | min_child_weight | 0-10 with increments of 1 |

## Notes

### Competing Interest Statement

All authors except of Maria-Anna Trapotsi and Andreas Bender are AstraZeneca employees

